# A Nanoheater-Integrated Fluorescence Lifetime Thermometer for Investigating Subcellular Heat Shock Factor 1 Responses

**DOI:** 10.64898/2026.09.02.748709

**Authors:** Hettimudalige Dilini Nisansala, Deshan Basnayake, Kayoko Nomura, Yohei Kono, Takeru Yamazaki, Yuya Matsuda, Cong Quang Vu, Takeshi Shimi, Satoshi Arai

**Affiliations:** Division of Nano Life Science, Graduate School of Frontier Science Initiative, Kanazawa University, Kakuma-machi, Kanazawa, 920-1192, Japan; WPI Nano Life Science Institute, Kanazawa University, Kakuma-machi, Kanazawa, 920-1192, Japan

## Abstract

Subcellular thermal engineering provides a powerful approach for investigating and manipulating biological processes. However, existing subcellular heating platforms capable of combining spatially confined heating, quantitative thermometry and simultaneous imaging of cellular responses remain limited. We developed a quantitative nanoheater-thermometer (qNanoHT), a polymeric nanoparticle integrating a temperature-sensitive fluorescent, dye and a photothermal dye. qNanoHT determines local temperature from fluorescence lifetime using fluorescence lifetime imaging microscopy (FLIM), thereby reducing susceptibility to photobleaching, focal drift and variations in probe concentration compared with intensity-based methods. The platform enabled real-time measurement at a subcellular heat spot while the dynamics of heat shock factor 1 (HSF1) were monitored in living cells. Heating at a single intracellular site was sufficient to induce HSF1 foci. Foci induced by mild heating at approximately 38 °C dissolved after heating ceased, whereas those induced by stronger heating at approximately 41 °C persisted and were associated with caspase-3/7 activation and apoptosis. Notably, qNanoHT-mediated subcellular heating induced HSF1 foci at a lower measured temperature than uniform whole-cell heating approximately 38 °C versus 39 °C indicating that the spatial extent of heating influences the HSF1 activation threshold. qNanoHT therefore provides a quantitative platform for relating local intracellular temperature to cellular stress responses and subsequent cell fate.

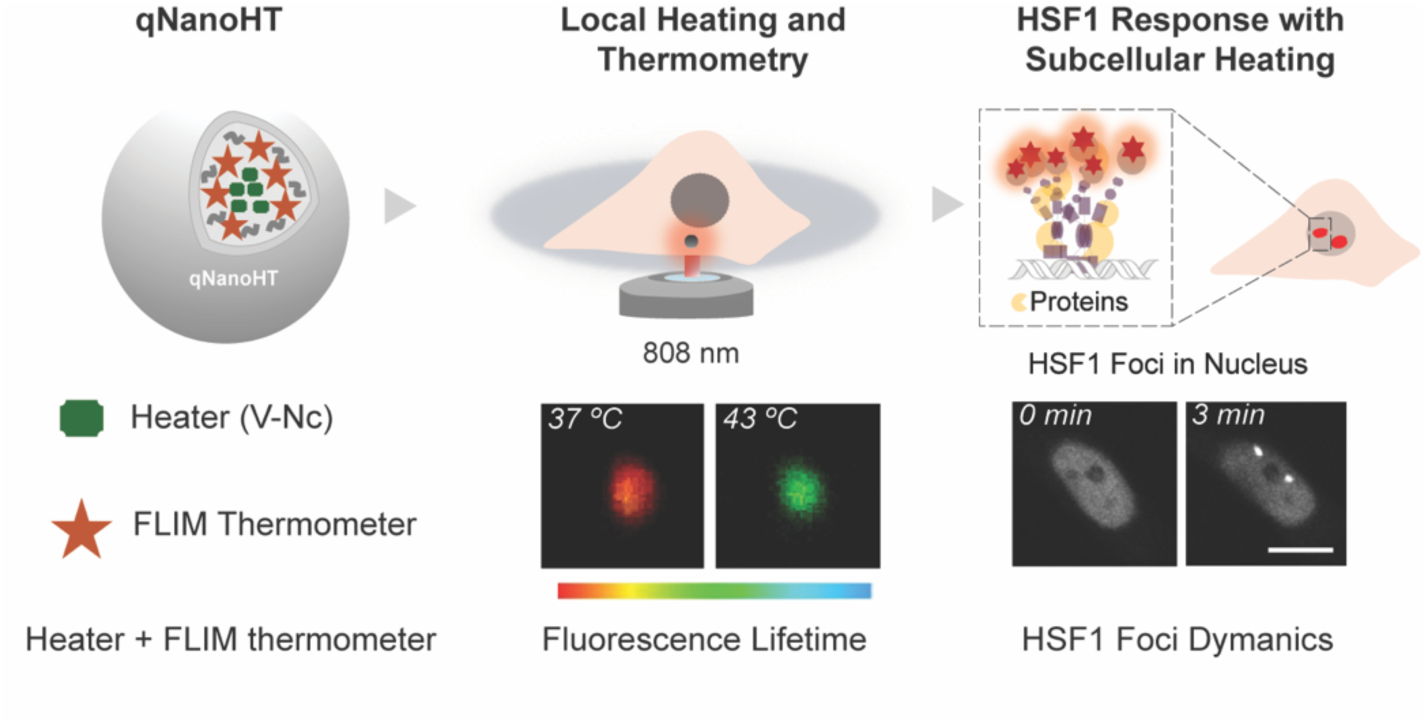

## Introduction

Thermal perturbation influences a wide range of cellular processes,^1^ including protein folding, metabolism,^2^ and signal transduction.^3–5^ Precise temperature modulation is therefore a powerful approach for elucidating the thermodynamic basis of these processes and manipulating cellular function in a controlled manner.^6^ Over recent decades, methods for controlling temperature in living cells have progressed from bulk heating towards increasingly localised techniques. Previous studies primarily relied on microscope stages or chamber-based heating systems, which uniformly regulate temperature across the entire sample. However, the slow thermal equilibration of these systems limits their ability to resolve dynamic cellular processes. To address this limitation, optical heating strategies have been explored, offering superior spatiotemporal precision.^78^ For example, optical microscopic heating systems that employ near-infrared (NIR) lasers (e.g., at 1470 nm) generate microscale heated regions through the vibrational excitation of water molecules.^9^ Nevertheless, their spatial resolution is constrained by the size of the laser focus, resulting in microscale heating that affects both the target region and surrounding cellular components. To overcome this constrain, nanosizd photothermal agents, including plasmonic metal nanoparticles^10^ and organic NIR dyes,^11,12^ have been developed to concentrate light absorption and heat generation within a nanoscale volumes. Because heat is generated directly by the nanosized agent, these material confine heating to a substantially smaller region than can be achieved using laser irradiation alone, thereby improving spatial resolution.

Localised heating provides a powerful strategy for dissecting local cellular responses at the subcellular level.^6,7^ However, precise and quantitative thermal modulation requires simultaneous thermometry during heating, with the thermometer positioned close to the heat source to accurately measure the local temperature.^7^ Integrating heating and thermometry functionalities within a single platform has therefore attracted considerable interest. Several hybrid heater–thermometer systems have been reported, pairing inorganic thermometric nanomaterials such as diamond particles,^13,14^ and quantum dots.^15^ However, these inorganic materials are typically bulkier and raise concerns regarding cytotoxicity, which can limit their suitability for minimally invasive intracellular applications.^16,17^ In the present study, we instead selected organic dyes for both heating and thermometry, as their smaller size and lower toxicity make them better suited for co-encapsulation within a single polymeric nanoparticle. Their biocompatibility and biodegradability may further reduce the invasiveness of intracellular applications.^18,19^ Based on this concept, we previously developed a polymeric nanoheater–thermometer that generates a subcellular heat spot within living cells under NIR irradiation.^20^ To improve thermometric accuracy, we incorporated a temperature-insensitive reference dye alongside a fluorescent thermometer, thereby enabling ratiometric fluorescence thermometry. Nevertheless, differences in the photobleaching rates of the two dyes can alter the fluorescence ratio over time, producing temperature independent changes that compromise measurement accuracy.

In contrast, fluorescence lifetime is an intrinsic property of the excited-state decay process and is therefore independent of probe concentration, excitation intensity, and photobleaching, providing a readout that avoids the artifacts common to intensity-based measurements.^21^ We therefore incorporated a fluorescence lifetime-base thermometer into our previously developed nanoparticle for quantitative temperature sensing at the heat spot.

In this study, we engineered fluorescence lifetime-based polymeric nanoparticles that integrate a photothermal heating dye and a temperature-sensitive dye; we, termed these nanoparticles quantitative nanoheater-thermometers (qNanoHTs). Temperature is estimated from the fluorescence lifetime of the temperature-sensitive dye using fluorescence lifetime imaging microscopy (FLIM), rather than from its fluorescence intensity. For this purpose, we employed a 4,4-Difluoro-4-bora-3a,4a-diaza-s-indacene (BODIPY) derivative as the fluorescent thermometer.^22^

To demonstrate the utility of the FLIM-based qNanoHT system for investigating cellular processes in living cells, we examined the dynamics of heat shock factor 1 (HSF1). HSF1 is a key transcription factor and the master regulator of the heat shock response. Upon thermal stress, HSF1 becomes activated and accumulates at specific nuclear sites, forming foci, that reflect its activation state and drive the transcription of heat shock proteins to restore protein homeostasis.^23,24^ Monitoring HSF1 foci formation therefore provides a useful model for evaluating heat-induced cellular responses, yet how a single intracellular heat spot influences HSF1 foci dynamics remains largely unexplored. Using qNanoHT, we investigate how subcellular heating regulates HSF1 activation and nuclear foci formation in living cells. Furthermore, by monitoring caspase activity alongside HSF1 dynamics, we examined how the reversibility of HSF1 activation influences subsequent cell fate.

## Results and Discussion

### Preparation and characterisation of qNanoHT

We designed a polymeric nanoparticle with two integrated functions: temperature quantification through fluorescence lifetime imaging of the incorporated BODIPY dye, and photothermal heating through an NIR-absorbing dye. A key consideration in the design of fluorescent thermometers is the physicochemical behaviour of BODIPY as a molecular rotor.^25^ As a molecular rotor, BODIPY exhibits viscosity-dependent fluorescence, arising from the coupling between local molecular mobility and the intramolecular motion of the BODIPY derivative in its excited state.

We therefore hypothesized that changes in temperature would alter the segmental mobility and effective local viscosity of the polymer matrix, thereby affecting the intramolecular motion and non-radiative relaxation of the embedded BODIPY. Consequently, these temperature-induced changes in the local polymer environment were expected to produce measurable changes in its fluorescence properties, enabling optical temperature sensing within the nanoparticle.

The viscosity sensitivity of BODIPY rotors is inversely correlated with the energy barrier associated with their transformation from a planar structure to a butterfly-like form.^26^ The unsymmetrical BODIPY derivative used in this study (Fig. 1A) was, developed by modifying the BODIPY core. It has been reported to exhibit relatively high viscosity sensitivity compared with the other BODIPY derivatives examined in the referenced study, rendering it a promising scaffold for the fluorescence-based detection of changes in local microviscosity.^22,27^

**Fig. 1.**
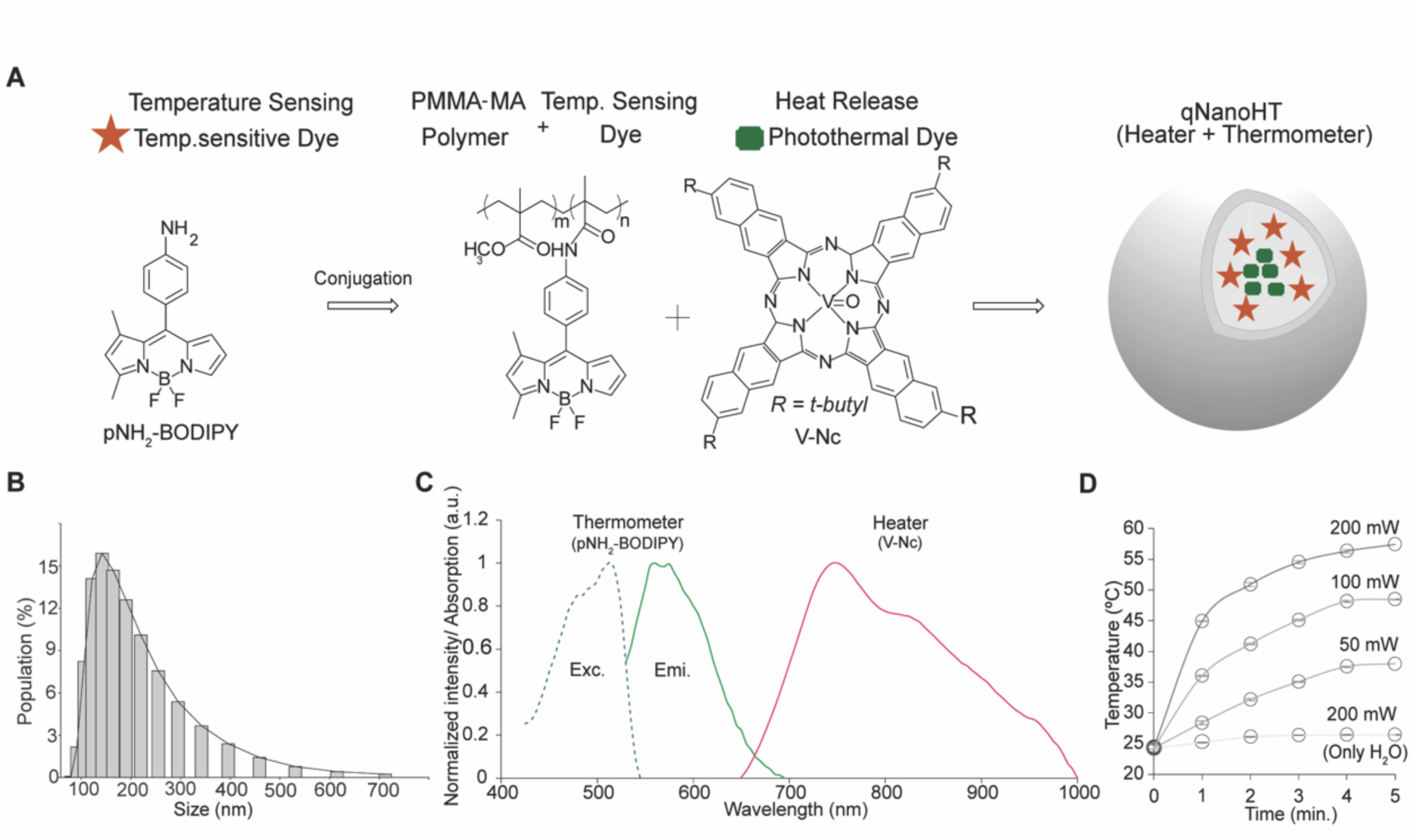
Preparation and characterisation of qNanoHT. (A) Schematic illustration of qNanoHT preparation by nanoprecipitation of PMMA-MA conjugated to amine functionalized BODIPY (pNH_2_-BODIPY) in the presence of the photothermal dye vanadyl 2,11,20,29-tetra-*tert*-butyl-2,3-naphthalocyanine(V-Nc). (B) Hydrodynamic size distribution of qNanoHT measured by dynamic light scattering (DLS). (C) Excitation and emission spectra of pNH_2_-BODIPY, and the absorption spectrum of V-Nc within qNanoHT. (D) Temperature increases in a qNanoHT suspension during irradiation with an 808 nm NIR laser. The initial temperature was 24 °C. Error bars represent the standard deviation, (n = 3).

First, we synthesized pNH_2_-BODIPY-conjugated PMMA-MA by covalently coupling pNH_2_-BODIPY to the polymer backbone (Fig. 1A). Because unconjugated BODIPY dyes can leak from polymeric nanoparticles after cellular uptake, covalent conjugation was used to retain the dye within the nanoparticle matrix.^28^ The resulting polymer was subsequently used as the matrix for nanoparticle preparation. To determine whether covalent conjugation prevented dye leakage, we compared nanoparticles containing physically encapsulated pNH_2_-BODIPY by fluorescence microscopy. Physically encapsulated pNH_2_-BODIPY produced diffuse intracellular fluorescence, indicating the leakage of the dye from the nanoparticles. By contrast, polymer-conjugated pNH_2_-BODIPY remained confined to discrete nanoparticle-associated puncta, suggesting that covalent conjugation effectively prevents intracellular dye leakage (Fig. S1).

As the photothermal agent, we selected V-Nc, an NIR-absorbing dye with strong absorption at 808 nm^29^ Owing to its high hydrophobicity, V-Nc was physically entrapped within the hydrophobic polymer core, without the need for covalent conjugation, and was incorporated into the pNH2-BODIPY-conjugated polymer nanoparticles by nanoprecipitation.^20,30^ The resulting nanoparticles had a hydrodynamic diameter of 135 ± 11 nm (Fig. 1B) and a zeta potential of 13 mV in cell culture medium. For our purpose, the negatively charged, unmodified nanoparticle surface was well-suited to this study. Because our cellular studies focused on investigating the effects of a single, spatially defined intracellular heat spot rather than on maximising uptake across the cell population. Passive uptake of individual nanoparticles was therefore sufficient for the single-spot heating experiments. Consequently, we omitted the additional cationic surface modifications commonly used to enhance cellular uptake through electrostatic interaction with negatively charged cell membrane.^31^

The conjugated pNH_2_-BODIPY within the polymer matrix exhibited excitation and an emission maximum at 515 and 560 nm, respectively, while V-Nc displayed broad absorption between approximately 650 and 950 nm (Fig. 1C). The limited spectral overlap between the two dyes allowed pNH_2_-BODIPY to be selectively excited for FLIM-based temperature measurement, while V-Nc was independently activated by NIR irradiation at approximately 808 nm to generate local heat. This spectral separation enabled heating and temperature sensing within the same nanoparticle with minimal optical crosstalk. The photothermal performance of qNanoHT was subsequently evaluated in an aqueous suspension (Fig. 1D). During irradiation with an 808 nm laser, the temperature of the qNanoHT suspension increased from an initial temperature of 24 °C in a laser power dependent manner, demonstrating efficient photothermal conversion.

We additionally, investigated whether NIR irradiation of qNanoHT generated reactive oxygen species (ROS). Oxidative stress can also induce HSF1 foci formation,^32^ therefore, the contribution of ROS-mediated effects had to be minimised when investigating HSF1 dynamics induced by heat stress. ROS generation was assessed in both qNanoHT suspensions and qNanoHT treated cells using indicator “2′,7′-dichlorodihydrofluorescein diacetate (H₂DCFDA). In solution, ROS generation by qNanoHT was compared with that of Indocyanine Green (ICG), a photothermal agent that can also be excited at 808 nm. The ICG concentration was adjusted to match the photothermal output of qNanoHT (Fig. S2). After 3 min of irradiation, qNanoHT generated negligible ROS relative to ICG, indicating that the observed HSF1 responses were predominantly induced by thermal, rather than oxidative stress.

### Characterisation of FLIM-Based thermometry in qNanoHT

To evaluate the feasibility of FLIM-based thermometry using the pNH_2_-BODIPY dye incorporated into qNanoHT. We examined how focal-plane displacement along the z-axis affected the fluorescence lifetime of pNH_2_-BODIPY. Because heating can alter cellular morphology, such changes could introduce artifacts into fluorescence-based thermometry. The fluorescence lifetime remained largely unchanged during focal displacement, whereas the fluorescence intensity fluctuated substantially (Fig. 2A). These findings highlight an important advantage of FLIM over intensity-based thermometry in intracellular heating studies: fluorescence lifetime is considerably less sensitive to focal-plane displacement, at least over a range of several micrometers, and can therefore provide more reliable temperature measurements in dynamic cellular environments.

**Fig. 2.**
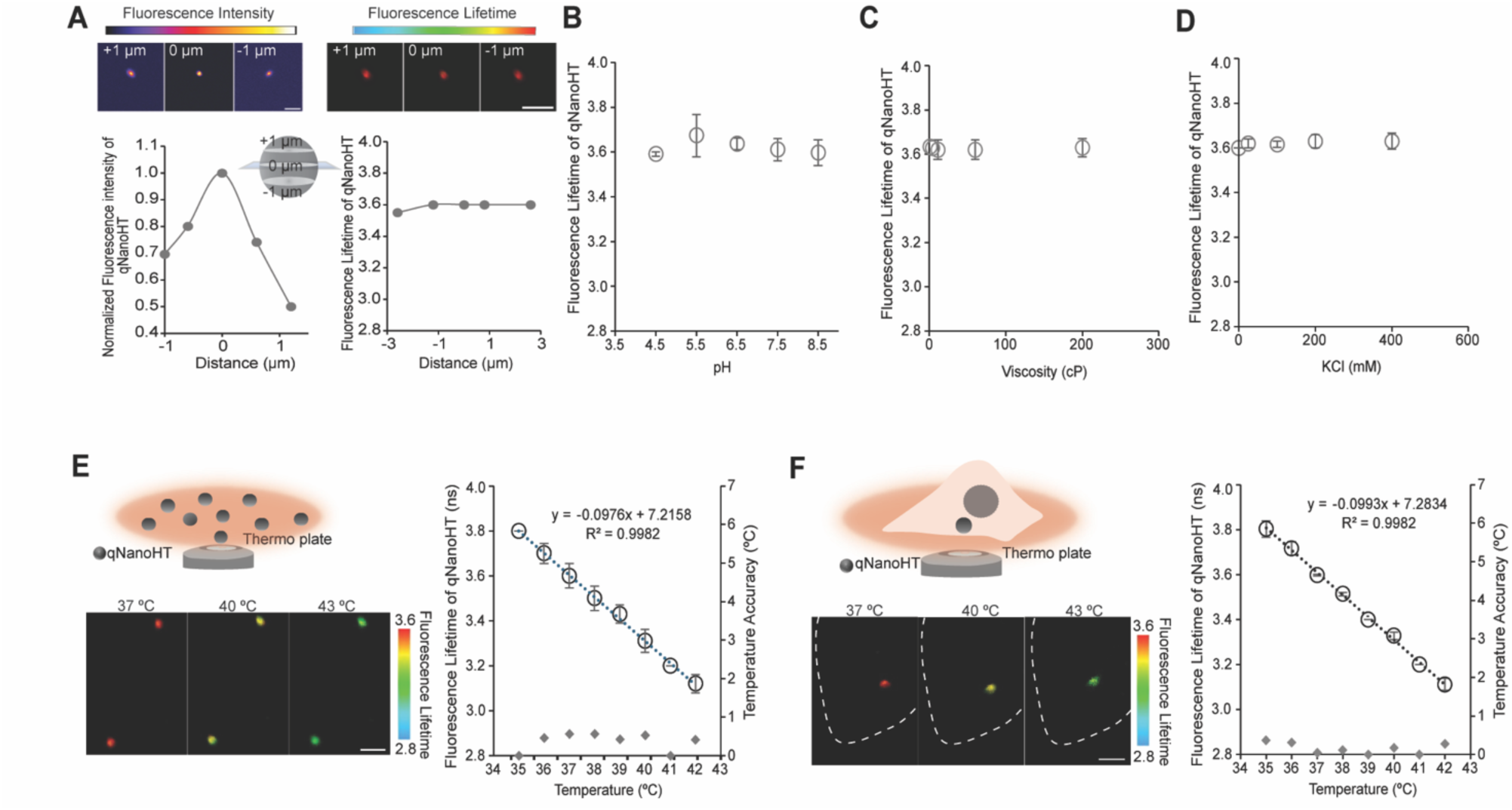
Characterisation and calibration of qNanoHT. (A) Effects of focal-plane displacement along the z-axis on the fluorescence intensity and lifetime of qNanoHT. (B-D) Robustness of the fluorescence lifetime of qNanoHT against variation in (B) pH (4.5–8.5), (C) viscosity (1–200 centipoise; cP), and (D) ionic strength (0–400 mM potassium chloride). Data are presented as the mean ± standard deviation, n = 3, (E, F) Temperature calibration of qNanoHT (E) on a glass-bottomed dish and (F) in living Hela-HSF1-Halo cells. Representative FLIM images and calibration curves are shown. The secondary axis indicates the accuracy of the temperature measurements.

We also evaluated whether the fluorescence lifetime was affected by pH (4.5–8.5; Fig. 2B), viscosity (1–200 cP; Fig. 2C) or, ionic strength (Fig. 2D). qNanoHTs internalised through endocytosis are typically retainedwithin acidic organelles such as lysosomes,^33^ consistent with the observed colocalization of qNanoHT woth lysosomal marker (Fig. S3), we assessed pH sensitivity over range encompassing the conditions expected within these acidic compartments. Sensitivity to external viscosity was evaluated using glycerol-water mixtures ranging from 1 to 200 cP. This range encompasses biologically relevant intracellular microviscosities, including those associated with crowded and condensed cellular environments. It was particularly relevant because intracellular viscosity can change during apoptosis,^34^ which was examined in the present study. Insensitivity to external viscosity is therefore important for maintaining accurate temperature measurement throughout the cellular response to heat stress. We further assessed the influence of ionic strength because intracellular compartments differ in their ionic compositions and local ion concentrations. The fluorescence lifetime exhibited negligible sensitivity to all three environmental factors (Fig. 2B–D). The negligible dependence on external viscosity is particularly noteworthy because the pNH_2_-BODIPY rotor is intrinsically designed to respond to changes in viscosity. If the dye were exposed at or near the nanoparticle surface, its fluorescence lifetime would be expected to respond to changes in the surrounding viscosity. The observed insensitivity therefore indicates that the pNH_2_-BODIPY was effectively embedded and shielded within the polymer matrix, rather than oriented toward the nanoparticle surface. Collectively, these results demonstrate that the fluorescence lifetime of qNanoHT is sufficiently robust for reliable intracellular thermometry.

We subsequently examined the temperature sensitivity of qNanoHT in acellular and intracellular environments. For dish-based measurements, a diluted qNanoHT suspension in cell culture medium was deposited onto a glass-bottom dish. For the intracellular experiments, cells were incubated with qNanoHTs for 12 h before experiment. This incubation period was selected based on cell viability assays showing negligible effects on cell viability (Fig. S4). The colocalization of qNanoHT with lysosomal markers was consistent with cellular uptake through the endocytotic pathway^35^ (Fig. S3). Fluorescence lifetime was measured across a range of temperatures using a microscope-mounted thermoplate. The temperature sensitivity of pNH_2_-BODIPY within qNanoHT was estimated to be 98 ps °C⁻¹ on the glass-bottomed dish and 99 ps °C⁻¹ in living cells (Fig. 2E, F). The mean temperature measurement accuracies were calculated as 0.39 ± 0.22 °C on the dish and 0.17 ± 0.13 °C in cells. These values are comparable to those reported for BODIPY-based fluorescent organelle thermometers, which range 0.45 ± 0.14 °C to 0.88 ± 0.27 °C).^21^ The PMMA matrix surrounding the pNH_2_-BODIPY does not appear to impede the intramolecular rotor motion required for, temperature-dependent changes in fluorescence lifetime. Moreover, qNanoHT exhibited comparable temperature sensitivities in the dish-based and intracellular environments. This consistency can be attributed to the embedding and shielding of pNH_2_-BODIPY within the polymer matrix, which prevents its direct exposure to the surrounding medium. Such shielding may explain why the temperature response of qNanoHT remained stable across the two environments, unlike those of organelle-targeted thermometers, whose direct exposure to distinct organelle microenvironments may result in different sensitivities.^21^

We further investigated the photothermal performance of qNanoHT using an 808 nm laser, corresponding to the absorption band of V-Nc. The temperature increase at the heat spot was quantified using the fluorescence lifetime–temperature calibration curves shown in Fig. 2E and F. A diluted qNanoHT suspension was deposited onto a glass-bottomed dish, and a single qNanoHT particle was irradiated with the 808 nm laser at powers ranging from 10 to 50 mW. The fluorescence lifetime decreased in a power-dependent manner, consistent with an increase in temperature. By contrast, qNanoHTs outside the irradiated region exhibited negligible changes in fluorescence lifetime (Fig. 3A), indicating that heat generation was spatially confined to the laser irradiated qNanoHT. At a given laser power, the magnitude of the temperature increase varied among qNanoHTs, potentially because of differences in the amount of V-Nc loaded into each nanoparticle. During irradiation, the fluorescence lifetime decreased and then returned to its initial value when the laser was switched off, demonstrating the reversibility of the thermal response (Fig. S5).

**Fig. 3.**
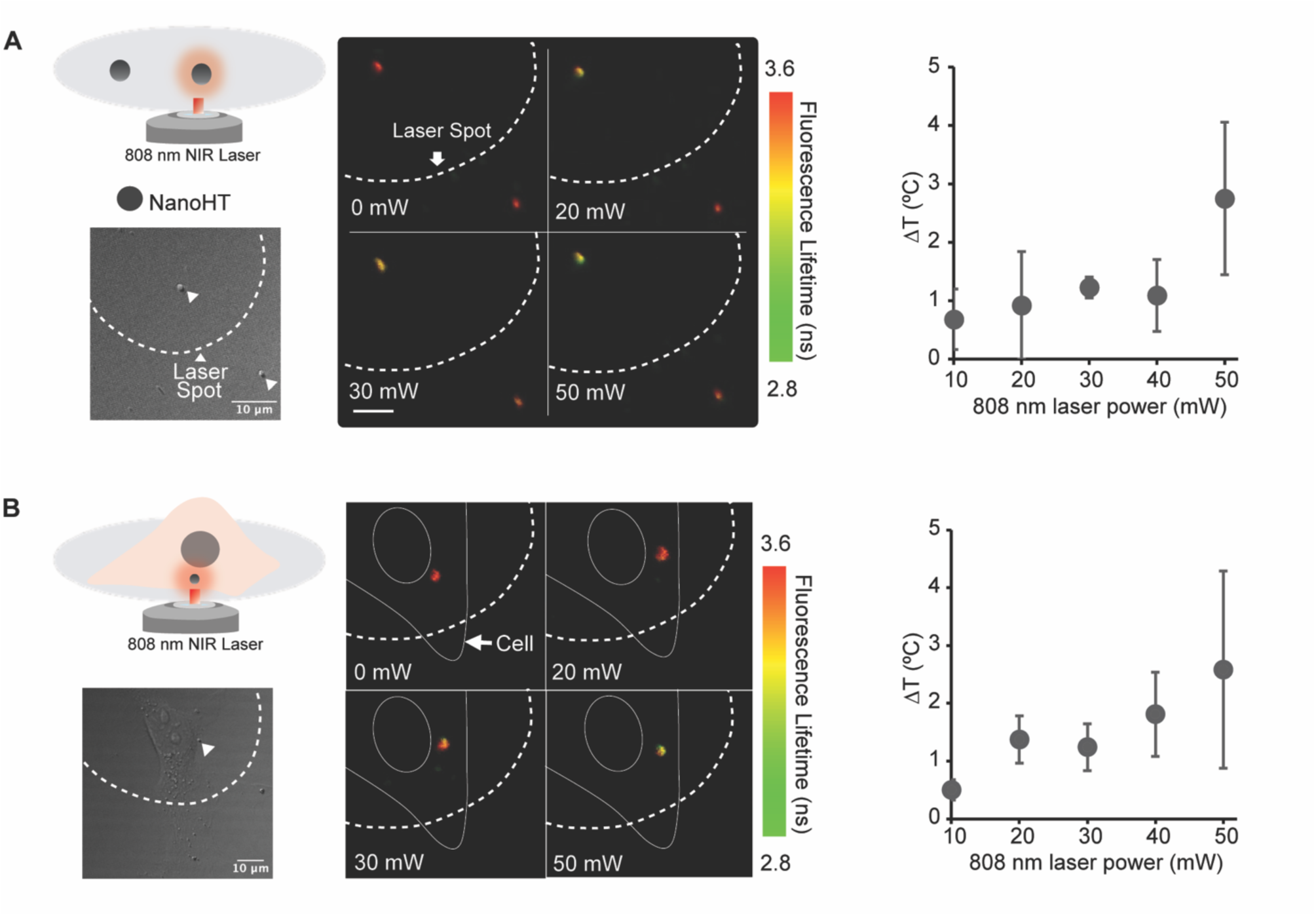
Validation of heat generation by qNanoHT during irradiation with an 808 nm NIR laser. (A) On a glass-bottomed dish and (B) within Hela-HSF1-Halo cells. The baseline temperature was 37 °C.

We subsequently evaluated the photothermal performance of qNanoHT following uptake by HeLa-HSF1-Halo cells. Consistent with the dish-based results, intracellular qNanoHTs generated comparable temperature increases irradiation with the 808 nm laser (Fig. 3B). The measured temperatures were within the biologically relevant mild hyperthermia (39-42 °C). This range is generally below the temperature used for ablative photothermal therapy and is therefore less likely to induce extensive necrotic cell death.^36^

We used qNanoHT, to investigate the heat shock response, with particular emphasis on the dynamics of HSF1. Under, heat stress, HSF1 is released from chaperone complexes and translocates to the nucleus, where it forms condensates known as nuclear foci.^37–39^ These foci form rapidly following heat exposure and can therefore serve as a sensitive, real-time indicator of cellular thermal stress. We established a system for tracking HSF1 foci dynamics within the nucleus using HSF1 fused to HaloTag and labelled, with the JF646 ligand. To validate this system as a reporter of HSF1 foci formation following thermal stimulation, the entire cell was heated using a 1470 nm laser.

This approach enabled real-time monitoring of heat-induced HSF1 formation.^40^ Temperature changes during irradiation with the 1470 nm were monitored using an endoplasmic reticulum (ER)-targeted fluorescent thermometer, ER Thermo Yellow (ETY) which has a temperature sensitivity of - 3.9% °C⁻¹.^41^ This thermometer was selected because the ER is distributed throughout the cell, allowing temperature changes to be measured across the heated region (Fig. 4A). Using the reported calibration, changes in ETY fluorescence intensity were converted into temperature increase (ΔT). HSF1 foci formation required a threshold temperature of approximately 39 °C. Moreover, at least 3 min of heating was required before the foci area reached a plateau at each temperature tested (Fig. S6).

**Fig. 4.**
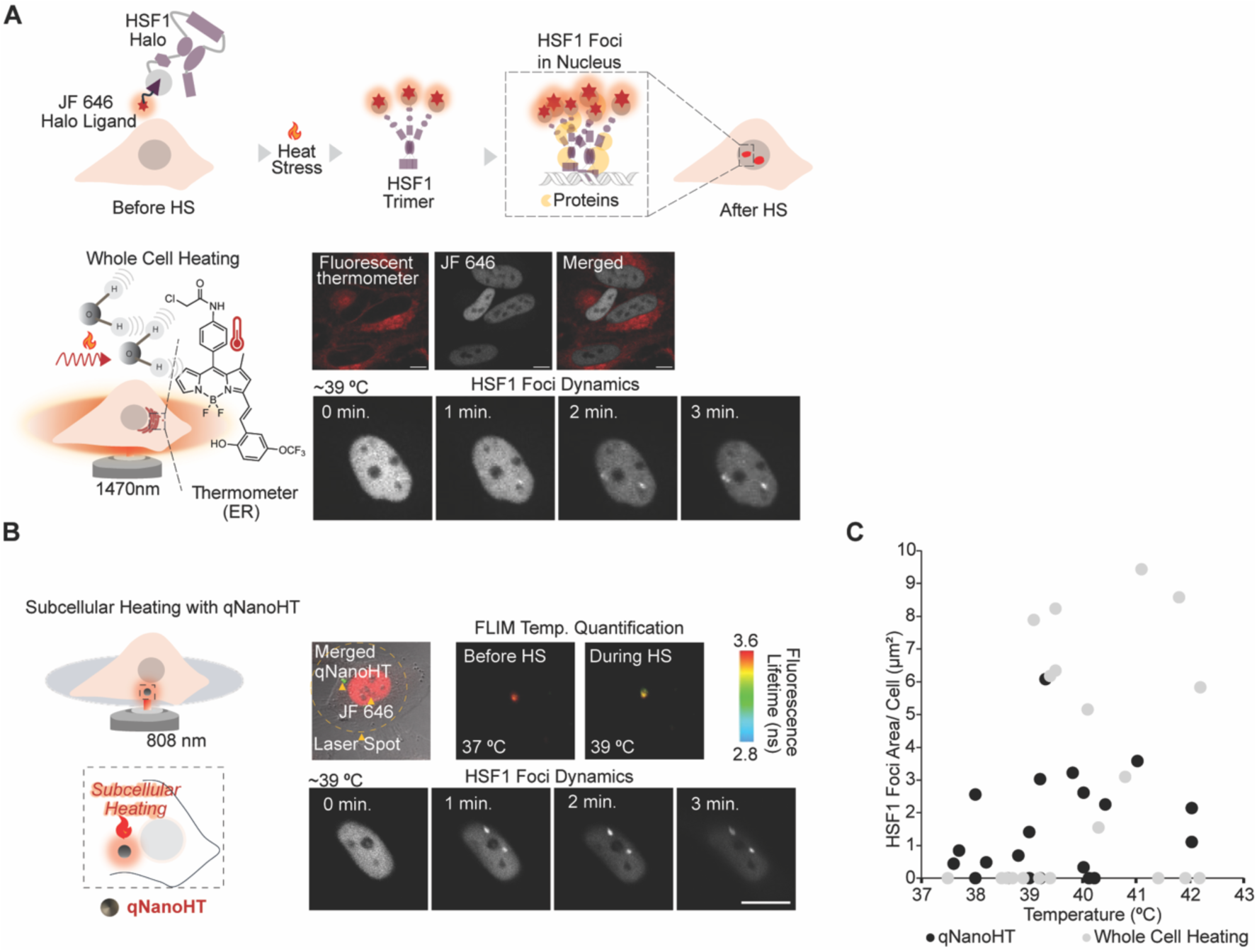
HSF1 foci formation following heat stress in HeLa cells expressing HSF1-HaloTag labelled with JF 646. (A) Whole-cell heating using 1470 nm laser. (B) Subcellular heating using qNanoHT and an 808 nm laser. (C) Comparative analysis of the HSF1 foci area following whole-cell and qNanoHT -mediated subcellular heating.

Having established this microscopic platform, we investigated whether localised heating at a single intracellular site was sufficient initiate the heat-shock response. Whereas irradiation with the 1470 nm laser heated the entire cell, qNanoHT combined with an 808 nm laser enabled spatially confined heating. The 808 nm laser power was limited to 50 mW because higher powers caused bulk heating of the medium through water absorption. Localised heating at a single spot was sufficient to induce HSF1 foci formation within the nucleus (Fig. 4B). This result demonstrated that qNanoHT-mediated heating is sufficient to activate the heat shock response without whole-cell heating, highlighting the sensitivity of HSF1 to local thermal perturbations. Whole-cell heating generally required temperatures above approximately 39 °C to induce HSF1 foci, whereas qNanoHT-mediated subcellular heating induced foci at approximately 38 °C (Fig. 4C). The difference between these thresholds exceeded the reported temperature measurement uncertainty of qNanoHT (0.17 ± 0.13 °C), suggesting that cellular responses differed between the two heating modes. We also observed variability in the number of nuclear foci, possibly reflecting inherent heterogeneity within the HeLa cell population. Given the reported involvement of HSF1 in cancer cell division, HSF1 foci formation may also vary according to the stage of cell-cycle.^42,43^ This possible relationship was not examined in the present study and warrants further investigation.

### HSF1 dynamics and apoptosis induced by subcellular heating

To understand the cellular consequences of HSF1 foci formation, we investigated whether qNanoHT-mediated heating activated apoptotic signaling and how this process was related to HSF1 dynamics. HSF1 foci formation is closely associated with cell-fate decisions under heat stress. Transient, liquid-like foci indicate an active heat-shock response; they form through liquid–liquid phase separation and promote chaperone gene expression.^44^ Under persistent stress, these condensates undergo a transition to a gel-like state associated with reduced chaperone induction and increased susceptibility to apoptosis.^45^. To assess the relationship between HSF1 foci dynamics and apoptotic signaling, we performed dual live-cell imaging of cells expressing labelled HSF1 together with either a caspase-8 or caspase-3/7 activity reporter following qNanoHT-mediated heating. Caspase-8 and caspase-3/7 report distinct, sequential stages of the apoptotic cascade: initiation and execution, respectively.^46–48^ The caspase-3/7 probe is non-fluorescent at baseline and becomes fluorescent following caspase-3/7-mediated cleavage,^49^ whereas the caspase-8 reporter is expressed in the cytoplasm and translocates to the nucleus upon caspase-8 activation.^50^ Both reporters were validated using 2 µM staurosporine, which produced the expected caspase-3/7 fluorescence and nuclear translocation of the caspase-8 reporter (Fig. 5 and Fig. S7).

**Fig. 5.**
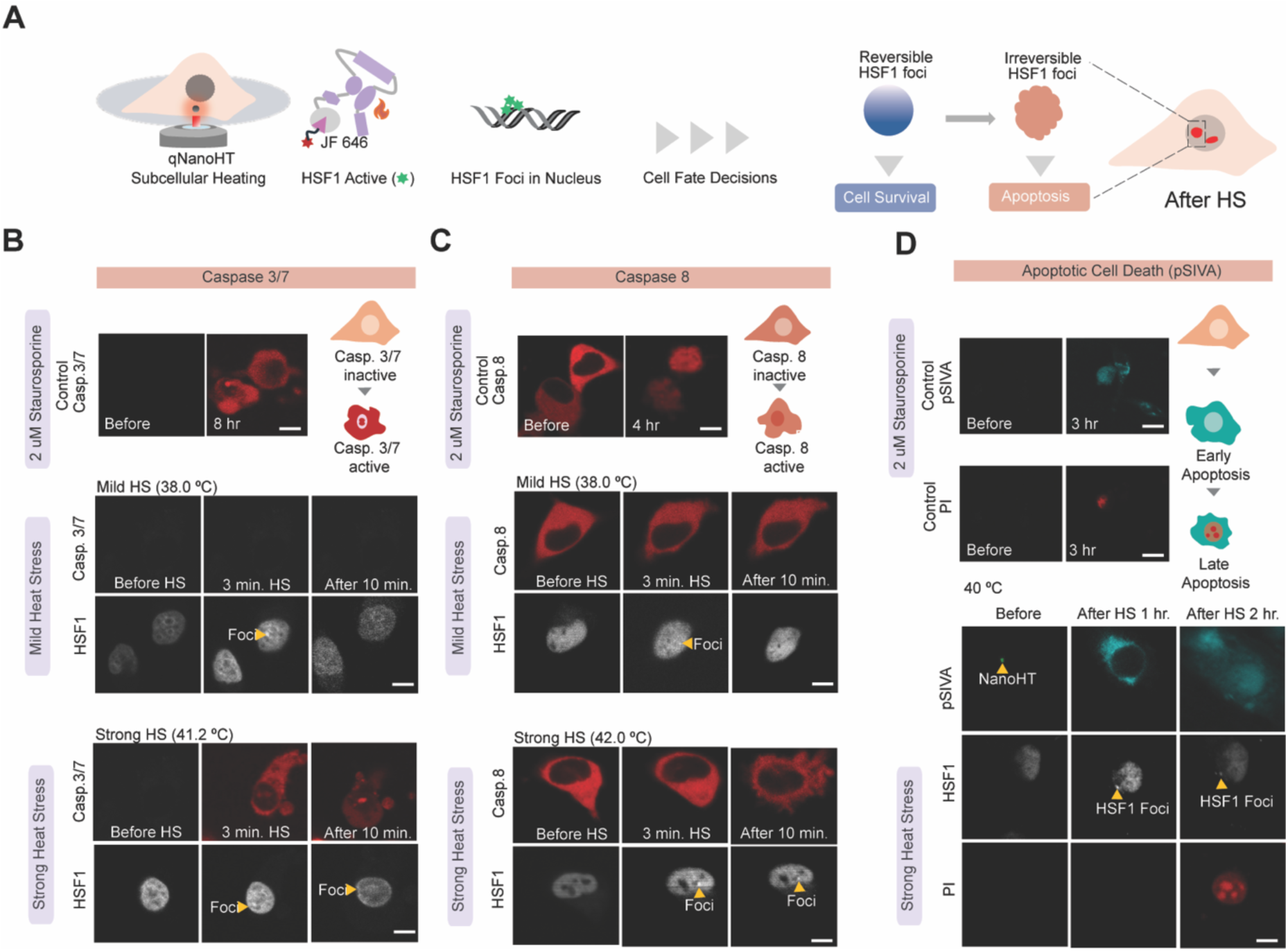
Cell death and HSF1 foci formation following subcellular heating. (A) Schematic illustration of HSF1 foci formation induced by subcellular heating, in which reversible foci are associated with cell survival, whereas irreversible foci are associated with apoptosis. Representative images of (B) caspase 3/7 activation and (C) caspase-8 activation together with HSF1 foci formation under mild and strong heat-stress conditions. (D) Polarity-sensitive indicator of viability and apoptosis (pSIVA) and propidium iodide (PI) labelling following qNanoHT-mediated cell death under strong heat stress.

Localised heating was applied using qNanoHT for 3 min and, was then discontinued. Under mild heat stress at approximately 38 °C, HSF1 foci formed but dissolved after the temperature returned to the baseline value, without subsequent caspase-3/7 activation. By contrast, under strong heat stress at approximately 41.2 °C, HSF1 foci persisted and rarely dissolved after heating was discontinued and the temperature returned to the baseline value of 37 °C. Caspase-3/7 activation was observed under these conditions (Fig. 5B). This finding is consistent with previous studies showing that persistent, irreversible HSF1 foci, rather than foci formation per se, serve as an indicator of apoptosis initiation.^51–54^ Despite clear caspase-3/7 activation following strong qNanoHT-mediated local heating no significant caspase-8 activation was detected (Fig. 5C). This observation is consistent with reports that caspase-8 is dispensable for heat-induced apoptosis, which instead proceeds through caspase-9-dependent, mitochondria signaling.^55,56^ The results therefore suggest that qNanoHT-mediated heat stress predominantly induces apoptosis through the intrinsic rather than the extrinsic pathway.^47,57^ Following the caspase experiments, pSIVA and PI assays were performed ^58^ to further assess whether qNanoHT-mediated cell death followed an apoptotic pathway. After a subcellular region was heated to 40 °C for 3 min using qNanoHT, pSIVA labeling was detected 1 h later, indicating early-stage apoptosis. PI labeling was detected after 2 h, consistent later loss of plasma-membrane integrity (Fig. 5D).

After characterizing the cell-death pathway, we considered the biological implications of spatially confined heating. A notable finding was that cellular responses to local heating differed from those observed during whole-cell heating (Fig. 4C). Specifically, the temperature thresholds associated with HSF1 foci formation and dynamics differed between the two heating modes. These observations suggest that heat-shock responses depend not only on the maximum temperature reached but also on the spatial extent of heating. Although qNanoHT is not targeted to a specific organelle, it is internalised through the endocytotic pathway and subsequently localizes to lysosomes. qNanoHT is therefore likely to generate heat primarily at or near lysosomes, producing a steep local temperature gradient similar to that previously characterised for NanoHT-mediated subcellular heating.^20^ By contrast, whole-cell heating exposes multiple organelles to heat simultaneously and relatively, uniformly. One possible explanation for the lower HSF1 activation threshold during subcellular heating is that local heating with qNanoHT selectively disrupt lysosomal homeostasis and autophagic flux, thereby generating a concentrated stress signal that efficiently activates HSF1.^59^ Whole-cell heating, in contrast, may activate protective mechanisms across multiple cellular compartments. For example, mitochondria and ER possess distinct stress-response pathways, in addition to the cytosolic response involving HSF1.^60^ These compartment-specific responses may mitigate cellular stress and alter the threshold for HSF1 activation. Although this interpretation remains speculative, it may partly explain the different activation thresholds observed under localised and whole-cell heating.

Alternatively, from a physical perspective, the local temperature gradient itself, rather than the measured local temperature alone, may contribute to HSF1 activation.^61,62^ Cellular temperature-sensing mechanism may be particularly sensitive to steep temperature gradient, potentially triggering HSF1 activation near heat-sensitive organelles even when the bulk cytoplasmic temperature remains relatively low. Although this interpretation is also speculative, it may help explain why localized heating induced HSF1 foci at a lower nominal temperature of approximately 38 °C than whole-cell heating.

In addition to activating HSF1, local heating with qNanoHT induced apoptosis. Caspase-3/7 activation in the absence of detectable caspase-8 activation suggests, that qNanoHT-induced heat stress predominantly activates the intrinsic, rather than the extrinsic, apoptotic pathway. However, further experiments assessing caspase-9 activation and cytochrome c release are required to confirm this interpretation and exclude a contribution from extrinsic signaling. Furthermore, qNanoHT is not targeted to a specific organelle, future studies should identify the precise subcellular site, and particularly the organelle, at which heat generation initiates HSF1 activation.

## Conclusions

We developed qNanoHT, an integrated a nanoheater-thermometer that enables simultaneous localised intracellular heating and fluorescence-lifetime-based temperature measurement with high spatial precision. qNanoHT-mediated subcellular heating induced nuclear HSF1 foci at a lower local temperature that whole-cell heating, whereas persistent foci under stronger heat stress were associated with caspase-3/7 activation and apoptotic cell death. Thus, combining localised thermometry provides a useful platform for investigating the spatial regulation of cellular heat stress responses in living cells. This technology may help identify unexplored intracellular thermal sensing mechanisms involved in HSF1 activation. Previous, nanoscale thermometry research has revealed heterogeneous intracellular temperature distributions and localised hot spots, indicating that cells may experience nanoscale temperature gradients rather than a uniform temperature field.^62^ In future, targeted photothermal technologies may support the development of more precise therapeutic approaches by enabling the selective modulation of HSF1 dynamics and cell death pathways.

### Experimental

#### Materials

PMMA-MA Mw ∼ 34,000 gmol^−1,^ M_n_ ∼ 15,000 gmol^−1^, vanadyl 2,11,20,29-tetra-tert-butyl-2,3-naphthalocyanine (V-Nc) were purchased from Sigma-Aldrich. Amine-functionalised Bodipy (pNH_2_-Bodipy) was synthesised according to a reported procedure.^63^

### Cell culture

HeLa cells stably expressing HSF1-HaloTag (HeLa-HSF1-Halo)^40^ were cultured in 3.5 cm glass-bottomed dishes containing Dulbecco’s modified Eagle’s medium (DMEM) supplemented with 10% fetal bovine serum and 1% penicillin-streptomycin. The cells were maintained at 37 °C in a humidified atmosphere containing 5% CO_2_.

### Preparation and characterisation of qNanoHT

PMMA-MA copolymer (M_n_ ∼ 15,000 gmol^−1^, 5.0 mg) was dissolved in anhydrous tetrahydrofuran (THF;800 µL), and after which 4-(4,6-dimethoxy-1,3,5-triazin-2-yl)-4-methylmorpholinium chloride (DMTMM;0.615 mg) was added. pNH_2_-BODIPY (1.0 mg) was dissolved separately in THF (200 µL) and added to the polymer solution. The reaction mixture was stirred overnight at room temperature. The pNH_2_-BODIPY-conjugated polymer was purified by precipitation in diethyl ether (20 mL), followed by centrifugation at 6000 rpm for 5 min. The resulting pellet was redissolved in THF (800 µL), after which V–Nc (0.8 mg) dissolved in THF (200 µL) was added. The mixture was slowly injected into Milli-Q water (5 mL) at a flow rate of 1 mL min⁻¹ under bath sonication to form the nanoparticles. THF was allowed to evaporate overnight in a fume hood. The resulting nanoparticles were purified twice using a PD-10 desalting column to remove unconjugated dyes.

### Evaluation of ROS Production

2’,7’-Dichlorodihydrofluorescein (H_2_DCF) was prepared by deacetylating 2’,7’-Dichlorodihydrofluorescein diacetate (H_2_DCFDA) as described previously.^64,65^ H_2_DCF was then used in an in vitro ROS-detection assay to evaluate ROS production by qNanoHTs. Briefly, 0.5 mL of 1.0 mM H_2_DCFDA in methanol was mixed with 2.0 mL of 0.01 M sodium hydroxide. The solution was incubated at 37 °C for 30 min to deacetylate H_2_DCFDA to H_2_DCF. The mixture was subsequently neutralized by adding 750 µL of 25 mM sodium dihydrogen phosphate, while monitoring the pH with a calibrated pH probe. The resulting non-fluorescent H_2_DCF solution was stored at −20 °C until use. All fluorescence measurements were performed in triplicate.

### Cell-viability assay

HeLa-HSF1-Halo cells were seeded in 96-well plates at a density of 5 × 10³ cells per well and cultured for 48 h at 37 °C in a humidified atmosphere containing 5% CO_2_ atmosphere. The culture medium was then removed, and 10 µL of the qNanoHT suspension was added at three dilution factors (1×, 10×, 100×). Deionised water (10 µL) was added to blank-control wells. Separate wells were incubated with qNanoHT for 4, 24, or 48 h. After each incubation period, the culture medium was replaced with medium containing 10% Cell Counting Kit-8 (CCK-8; Dojindo) reagent, and the cells were incubated for a further 4h under the same conditions. The plates were then centrifuged at 300xg for 3 min, and absorbance was measured at 450 nm using a microplate reader.

### Evaluation of the photothermal response of qNanoHT

The photothermal response of qNanoHT was evaluated as described previously.^66^ Briefly, an aqueous qNanoHT suspension (1 mL) in a quartz cuvette was irradiated with an 808 nm laser at powers of 50, 100, or 200 mW for 10 min. Water without qNanoHT was used as the control. Temperature was recorded at every 30 s interval using a TES 1310 Type-K thermocouple.

### Fluorescence imaging of qNanoHT

Confocal fluorescence images were acquired using an FV1200 confocal microscope (Olympus) equipped with a PLAPON 60× oil-immersion objective lens with a numerical aperture (NA) of 1.42. Images were acquired through the fluorescein isothiocyanate (FITC) channel using a 473 nm excitation laser and a 490–540 nm: emission filter. Each image comprised 512 × 512 pixels and was acquired 1.109 s per frame. Fluorescence-lifetime images were acquired using a rapidFLIM HiRes system equipped with a MultiHarp 150 time-correlated single-photon counting unit, (PicoQuant) a 485 nm pulsed laser and a 520/35 BrightLine HC emission filter. Images were images collected at 512 × 512 pixels and 1.109 s per frame and processed using SymPhoTime 64 software (PicoQuant). The instrument response function (IRF) was determined from the deviation between the measured and estimated fluorescence decay curves. The qNanoHT, fluorescence decay profile convolved with the IRF was fitted using a double-exponential decay model. Photothermal stimulation during imaging was performed using, an IR-LEGO-100/mini/E system (SIGMAKOKI) integrated with the microscope. This system enabled irradiation at 808 nm during image acquisition to excite V-Nc.

### Evaluation of the effects of pH, viscosity and ionic strength

To characterize the response of qNanoHT to pH, viscosity, and ionic strength, the following solutions were prepared: 10 mM acetate buffer at pH 4.5, 10 mM 2-(N-morpholino)ethanesulfonic acid (MES) buffer at pH 5.5 and 6.5, 1× phosphate-buffered saline (PBS) at pH 7.5, and 10 mM 4-(2-hydroxyethyl)-1-piperazineethanesulfonic acid (HEPES) buffer at pH 8.5. Viscosities of 0, 4, 11, 60, and 200 cP were prepared using glycerol-water mixtures at different ratios. Pottasium chloride concentrations of 0, 25, 100, 200, and 400 mM were used to evaluate the effect of ionic strength. qNanoHT was dispersed in each solution in a glass-bottom dish, and its fluorescence lifetime was measured using FLIM.

### Preparation of qNanoHT calibration curves

For dish-based calibration, qNanoHTs were dispersed in 2 mL of PBS and incubated overnight in 3.5 cm glass-bottomed dishes to allow the nanoparticles to settle onto the glass substrate. The temperature was varied from 34 to 42 °C using a microscope-mounted thermo-plate, and the fluorescence lifetime was measured at each temperature. For intracellular calibration, HeLa-HSF1-Halo cells were incubated with qNanoHTs overnight at 37 °C in an atmosphere containing 5% CO_2_. Before imaging, the cells were washed with PBS and transferred to phenol-red-free DMEM. The temperature was varied from 34 °C to 42 °C using the microscope-mounted thermoplate, and the intracellular fluorescence lifetime of qNanoHT was measured at each temperature.

#### Generation of HeLa-HSF1-HaloTag cells

For piggyBac transposon system-mediated stable introduction of the HSF1-Halo into HeLa cells, we followed the methods described previously.^67,68^ Briefly, 4 µg PB533-HSF1-Halo and 1 µg pcDNA3-mPB were mixed with 15 µL FuGENE4K (Promega) added on to a 100 mm dish. Transfected cells were selected by incubation in medium containing 400 µg/mL G-418 disulfate aqueous solution (Nacalai Tesque). Four days after positive selection, HeLa-HSF1-Halo cells were trypsinized and incubated with 100 nM Janelia Fluor 646 HaloTag Ligand (Promega) for 45 min, then HSF1-Halo-positive population was collected using the SH800S cell sorter (Sony Biotechnology).

#### Whole-cell and subcellular heating

HeLa-HSF1-HaloTag cells were incubated with qNanoHTs for 12 h at a concentration adjusted to achieve uptake of approximately one qNanoHT particle per cell. Before imaging, the cells were labelled with 50 nM Janelia Fluor 646 (JF646) HaloTag ligand dye (Promega) for 30 min. Two heating modalities were used, with the baseline temperature maintained at 37 °C using a microscope stage-top incubator. For whole-cell heating, a 1470 nm diode laser was directed across the entire cell. Temperature was measured using the endoplasmic reticulum (ER)-targeted fluorescent thermometer ER Thermo Yellow (ETY). Changes in ETY fluorescence intensity were converted into temperature increases using a calibration factor of -3.9% °C⁻¹. Temperature was measured after 20 s of heating, after which HSF1 foci were imaged. For subcellular heating, qNanoHT was irradiated with an 808 nm laser. The fluorescence lifetime of qNanoHT was recorded before and during 20 s of laser irradiation, and then the temperature increase was calculated using the intracellular calibration curve.

#### Analysis of HSF1 foci formation

Confocal microscopy was used to visualize HSF1 foci. Z-stack imaging was not performed to maintain consistency in the imaging plane and avoid changes in the photothermal heating profile caused by variations in focal depth. Preliminary experiments were performed to determine the minimum temperature and heating duration required induce HSF1 foci formation in HeLa HSF1-Halo cells. Based on the intracellular photothermal performance of qNanoHT, temperatures ranging from 38 to 42 °C were investigated. HSF1-HaloTag was labeled with JF646-HaloTag ligand (excitation: 635 nm; reflective mirror; emission filter: BA655-755). For whole-cell heating experiments, ETY was imaged using excitation at 559 nm, an SDM640 dichroic mirror and a BA575-620 emission filter. Images were analyzed using Fiji (imageJ). The mean HSF1 foci area per cell was calculated and plotted against temperature for both qNanoHT-mediated subcellular heating and whole-cell laser heating.

### Analysis of cell death in relation to HSF1 dynamics

Cells were incubated with 50 nM JF-646 HaloTag ligand for 30 min at 37 °C before imaging. Caspase-3/7 activation was monitored using the CellEvent caspase-3/7 Red Detection Reagent, with which the cells were incubated for 30 min before imaging. Caspase-8 activity was monitored using a genetically encoded fluorescent protein biosensor. ^50^ qNanoHTs were added to the cells and incubated for 12 h before imaging. During live-cell imaging, the nanoparticles were irradiated with an 808 nm laser at 50 mW for 3 min to generate localised heat spots. Temperature was quantified from the fluorescence lifetime of qNanoHT. Live-cell imaging was performed using an Olympus IX83 inverted microscope equipped with an FV12-FD detector and a PLAPON 60 × oil-immersion objective lens with an NA of 1.42. A stage-top incubator maintained the cells at 37 °C throughout the experiment. Images were acquired at 512 × 512 pixels. HSF1 foci were imaged using excitation at 635 nm and a BA655–755emission filter. Caspase-8 activity, reported by mCherry, and caspase-3/7 activity were examined in separate experiments using excitation at 559 nm and a BA575–675 emission filter.

## Data availability

The data supporting this article have been included as part of the ESI.

## Author contributions

Conceptualization, S.A. and H.D.N.; methodology, H.D.N., D.S., K.N., Y.K., T.Y., Y.M., C.Q.V., and T.S.; writing, H.D.N. and S.A.; supervision, S.A. All authors have read and agreed to the published version of the manuscript.

## Conflicts of interest

There are no conflicts to declare.

## Acknowledgement

This work was supported by the WISE Program for Nano-Precision Medicine, Science, and Technology at Kanazawa University (MEXT); JSPS KAKENHI grant numbers 24K01307 (to S.A.) and JP202616267 (Grant-in-Aid for JSPS Fellows); JST FOREST Program (JPMJFR201E to S.A.); and the World Premier International Research Center Initiative (WPI), MEXT, Japan. The authors used ChatGPT (OpenAI) and Claude (Anthropic) solely for English language editing and to improve the clarity and readability of the manuscript. The authors reviewed, revised, and verified all AI-assisted suggestions and take full responsibility for the final content.

## Notes

### Competing Interest Statement

The authors have declared no competing interest.

