## Supplementary information for "A Nanoheater-Integrated Fluorescence Lifetime Thermometer for Investigating Subcellular Heat Shock Factor 1 Responses"

### Supporting Information

Fig.S1 qNanoHT Characterization — Page 02

Fig.S2 Evaluation of ROS generation — Page 02,03

Fig.S3 Colocalization test of qNanoHT — Page 03

Fig.S4 Cell Viability — Page 05

Fig.S5 Reversibility of FL of NanoHT after heating — Page 06

Fig.S6 HSF1 foci formation with whole cell heating — Page 07

Fig.S7 Positive controls of caspase 3/7 and caspase 8 sensors — Page 07

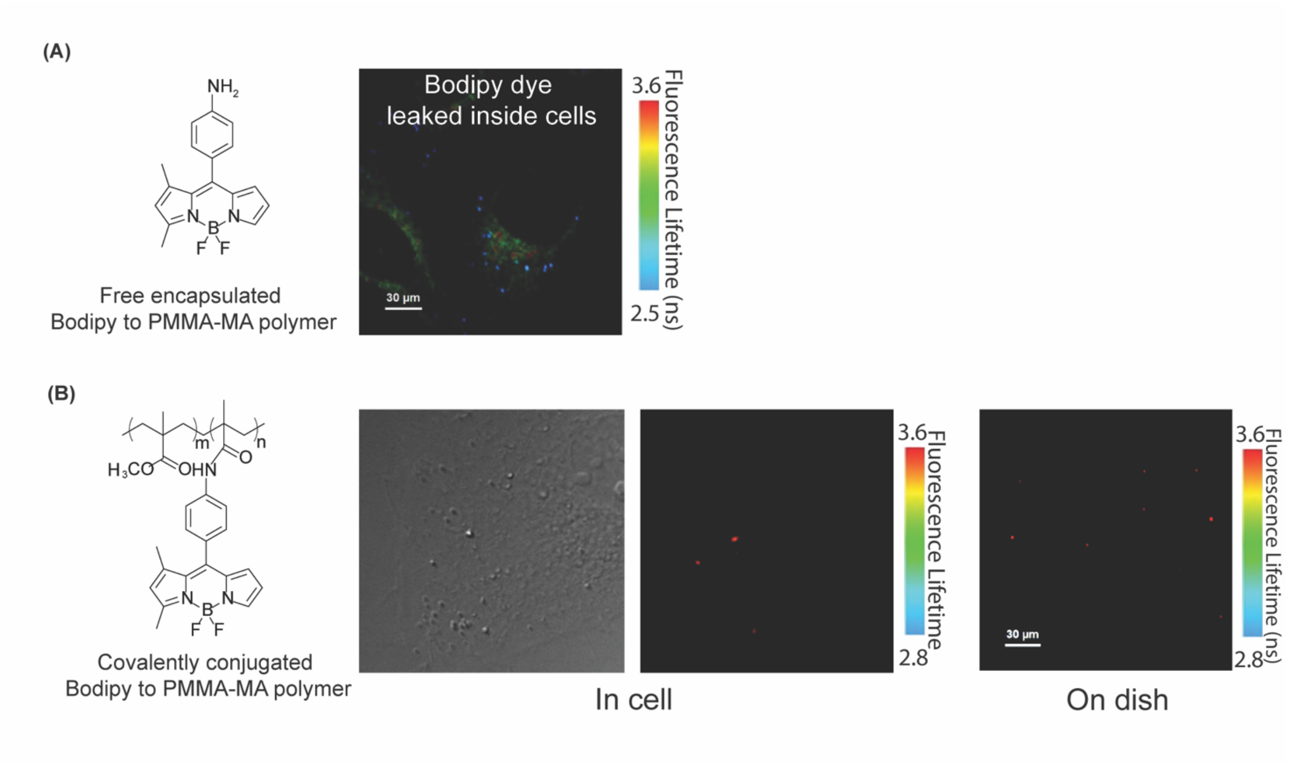


**Figure S1.** Dye leakage from qNanoHTs without and with BODIPY conjugation. Panel A: qNanoHTs containing non-conjugated BODIPY showed dye leakage. B) qNanoHTs containing conjugated BODIPY showed scarce detectable dye leakage.

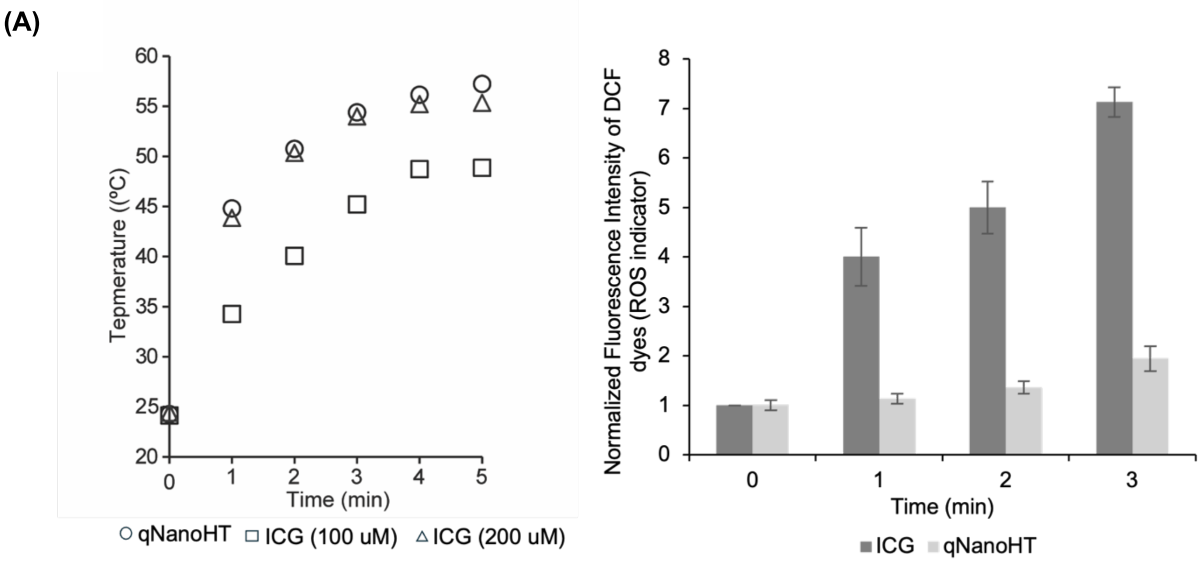

(B)

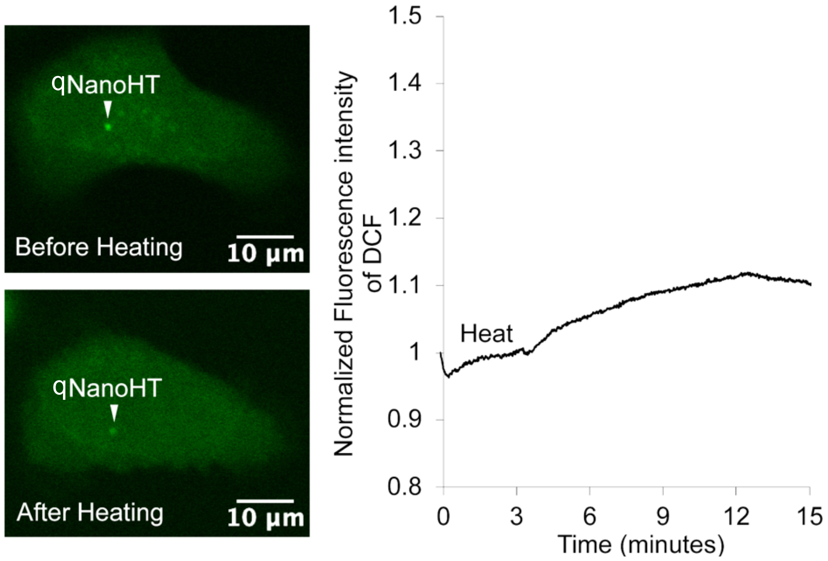


**Figure S2.** Evaluation of reactive oxygen species (ROS) generation by qNanoHT and ICG. (A) Photothermal heating curves of qNanoHT and 200 μM Indocyanine Green (ICG), showing comparable photothermal abilities. (B) ROS generation from qNanoHT and ICG in water following 808 nm laser irradiation (200 mW, 3 min). Error bars represent SD (n = 3). (B) Intracellular ROS imaging using a DCF dye.

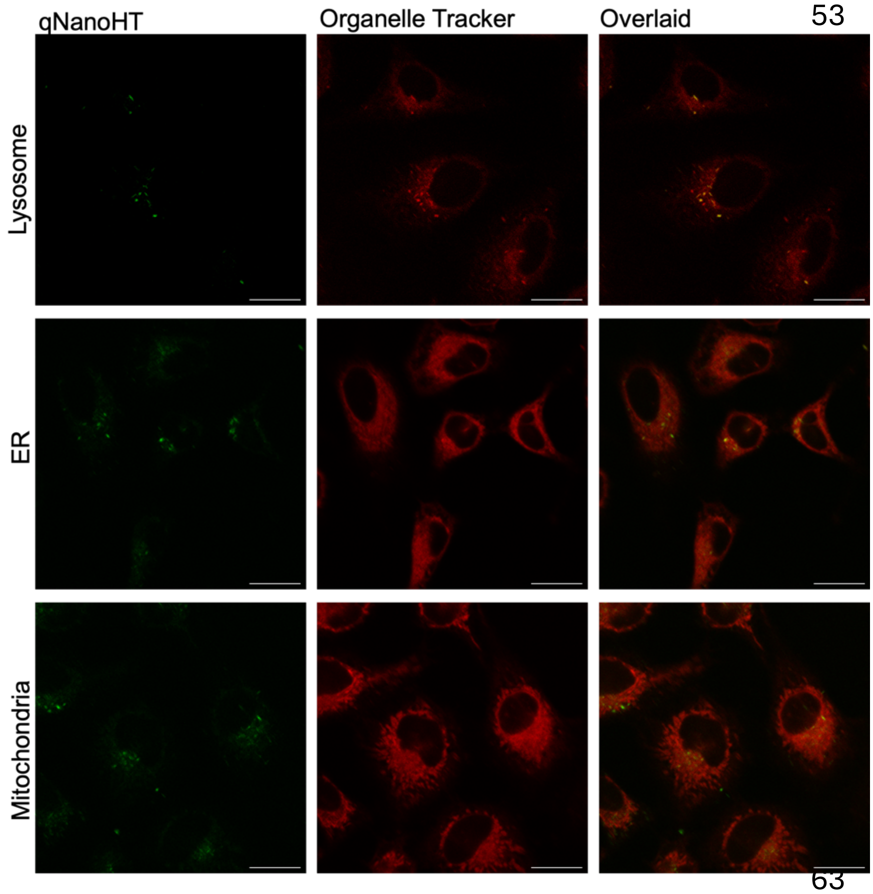


**Figure S3.** Colocalization test of qNanoHT after 12 hr incubation, scale bar 10 um.

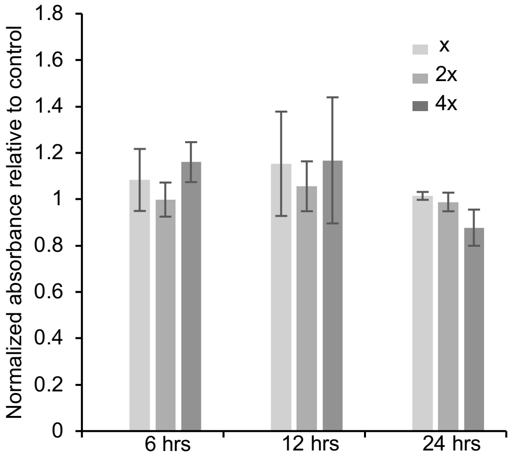


**Figure S4.** Cell viability evaluation of qNanoHT. Cell viability was assessed using a CCK-8 assay after treatment with three different concentrations of NanoHT (x, 2x, and 4x μg $mL^{-1}$). Error bars represent SD (n = 3). Cell viability did not decrease to 50% ($IC_{50}$) within the tested concentration range. Higher concentrations were not tested because qNanoHT is prone to aggregation in cell culture medium, likely due to interactions with serum proteins and salts; such aggregation can alter nanoparticle uptake and optical properties, or cause precipitation, and therefore would not represent a physiologically meaningful higher dose.

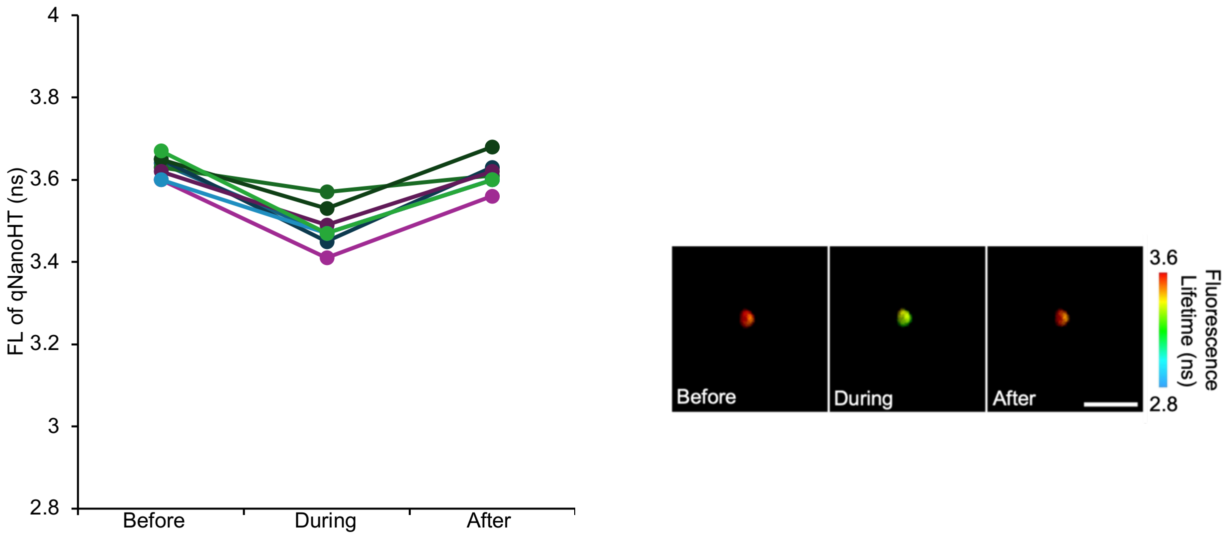


**Figure S5**. Reversibility of qNanoHT. qNanoHT in live cells were irradiated with an 808 nm laser (50 mW) at a baseline temperature of 37 °C. The fluorescence lifetime of qNanoHT returned to its initial value after irradiation, demonstrating reversible temperature responsiveness (n = 9).

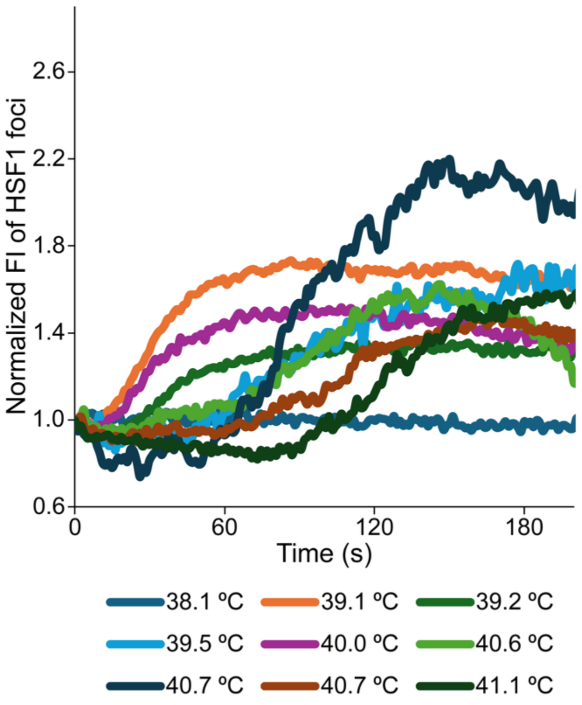


**Figure S6.** Increase in HSF1 foci fluorescence intensity after 3 min of whole-cell heating (by varying laser power (1470 nm).

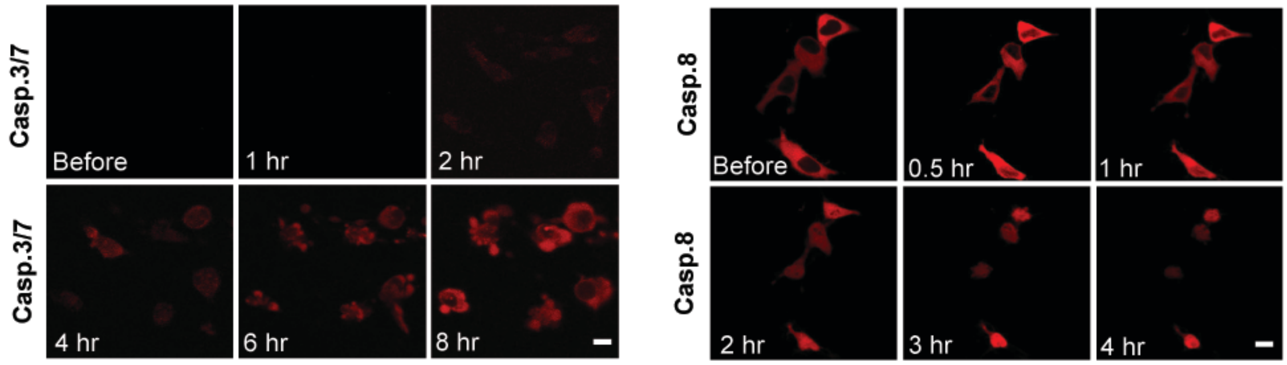


**Figure S7.** Time-lapse images of caspase 3/7 and caspase 8 sensors following treatment with 2 μM staurosporine.
